# A neural model for V1 that incorporates dendritic nonlinearities and back-propagating action potentials

**DOI:** 10.1101/2024.09.17.613420

**Authors:** Ilias Rentzeperis, Dario Prandi, Marcelo Bertalmío

## Abstract

The work of Hubel and Wiesel has been instrumental in shaping our understanding of V1, leading to modeling neural responses as cascades of linear and nonlinear processes in what is known as the “standard model” of vision. Under this formulation, however, some dendritic properties cannot be represented in a practical manner, while evidence from both experimental and theoretical work indicates that dendritic processes are an indispensable element of key neural behaviors. As a result, current V1 models fail to explain neural responses in a number of scenarios. In this work, we propose an implicit model for V1 that considers nonlinear dendritic integration and backpropagation of action potentials from the soma to the dendrites. Our model can be viewed as an extension of the standard model that minimizes an energy function, allows for a better conceptual understanding of neural processes, and explains several neurophysiological phenomena that have challenged classical approaches.

**Significance statement:** Most current approaches for modeling neural activity in V1 are data driven; their main goal is to obtain better predictions and are formally equivalent to a deep neural network (DNN). Aside from behaving like a black-box these models ignore a key property of biological neurons, namely, that they integrate their input via their dendrites in a highly nonlinear fashion that includes backpropagating action potentials (bAPs). Here, we propose a model based on dendritic mechanisms, which facilitates conceptual analysis and can explain a number of physiological results that challenge standard approaches. Our results suggest that the proposed model may provide a better understanding of neural processes and be considered as a contribution in the search of a consensus model for V1.

## Introduction

From V1 recordings in cat and primate, Hubel and Wiesel classified neurons into two types: simple cells, which respond maximally to oriented stimuli of a given contrast polarity and at a specific location, and complex cells, which show response invariance to position and polarity. Based on the neurons’ laminar locations and response properties, they proposed a hierarchical model (HM), where simple cells receive thalamic input, while complex cells receive input from simple cells (Hubel and Wiesel, 1962, 1968). Based on this conceptual model, the responses of the two types of cells were initially described mathematically in distinct ways: simple cells by a linear-nonlinear (LN) model where a linear filter is followed by a nonlinearity (Movshon et al., 1978; Heeger, 1992), and complex cells by an energy model where the outputs of phase-shifted filters are rectified and then added up (Adelson and Bergen, 1985). While the HM and the notion of simple and complex cells have been instrumental in shaping our understanding of V1, both concepts have been challenged by experimental studies that show the existence of complex cells that receive direct thalamic input (Martin and Whitteridge, 1984) and point to a continuum from simple to complex cell behaviors instead of a strict dichotomy (Rust et al., 2005; Chen et al., 2007). To account for these findings, several current models assume that V1 cells may receive both thalamic and cortical input and take the form of a cascade of LN elements (McFarland et al., 2013; Vintch et al., 2015).

These models attribute complex-like behavior to a larger contribution of cortical input, but complex cells can maintain their properties even when simple cells are silenced (Malpeli et al., 1986; Constantinople and Bruno, 2013), and some complex cells receive a strong monosynaptic drive from the thalamus (Su et al., 2024). For several experimental findings that contradict the simple/complex dichotomy, the explanations provided from within the standard model framework are not satisfactory. For instance, intracellular recordings in cat V1 show a change in the properties of the same cell depending on the type of stimulus, with a simple-like behavior for dense noise stimuli that switches to a more complex-like one for sparse noise stimuli (Fournier et al., 2011). Another challenging neurophysiological phenomenon is the increase in phase sensitivity of complex cells as stimulus contrast decreases (Crowder et al., 2007; Van Kleef et al., 2010): HM variations (Van Kleef et al., 2010) and recurrent amplification models (Yunzab et al., 2019a) cannot explain complex cell behavior for thalamic input; and data-driven models must adapt the number and structure of their filters and nonlinearities, to different inputs (Almasi et al., 2020).

Extensive evidence points to the importance of dendrites in neural processing (Pagkalos et al., 2024; Poirazi and Papoutsi, 2020). Notably, dendrites can receive backpropagating action potentials (bAPs) initiated from the soma (Stuart and Spruston, 2015) that can significantly influence neurons’ responses (Francioni and Harnett, 2022; Stuyt et al., 2022; Stingl et al., 2025). Since bAPs allow for a back and forth interaction between cell output and dendritic contributions, they cannot be represented in a practical manner under an LN formulation, requiring instead an approach based on differential equations (Larkum, 2022).

Here, we propose an extension of the classical formulation of the standard V1 model that considers both linear and nonlinear dendritic integration of the input, with the latter operating in a dynamic and input-dependent manner since processing is affected by bAPs. The proposed model produces a continuum from simple to complex cell responses, explains the emergence of complex cell responses for thalamic input, and the effect of stimulus configuration, statistics, and contrast on the response properties of cells. It is also robust to noise perturbations in its components, which is a prominent feature of biological systems (Marom and Marder, 2023; Albantakis et al., 2024). In our formulation, backpropagating responses contribute positively to the emergence of complex cell behavior and the sharpening of orientation-tuning curves.

## Materials and Methods

### Proposed model

We propose a model for V1 cells that can be considered an extension of the point neuron model, where in addition to a term corresponding to linear integration of inputs, as in the standard model, there are also two other terms corresponding to different types of dendritic nonlinearities, both including bAPs.

Our model equation of the equilibrium membrane potential *v*_*i*_ for cortical neuron *i* is

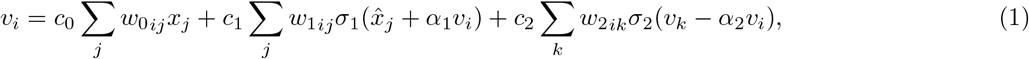

where each of the terms on the right represents a different dendritic process; *c*_{0,1,2}_ are scalar values that weigh the contribution of each term, acting as hyperparameters since they vary depending on the hypothesis we test; and *w*_{0,1,2}_ are weight matrices that act as filters for each of the dendritic processes.

The first term in Eq. 1 represents the filtering operation performed by linear dendrites (**Fig. 1a**). Here *x*_*j*_ is the firing rate input elicited by a static spatial pattern at position *j*. This input comes from a cell in an upstream layer, either thalamic or cortical, and has a plus sign for an ON-center input cell, and a minus sign for an OFF-center one. For no contribution from the nonlinear terms (*c*_1_, *c*_2_ = 0), the model is reduced to the point neuron model with the linear filter *w*_0_ denoting the classical receptive field of simple cells.

**Figure 1:**
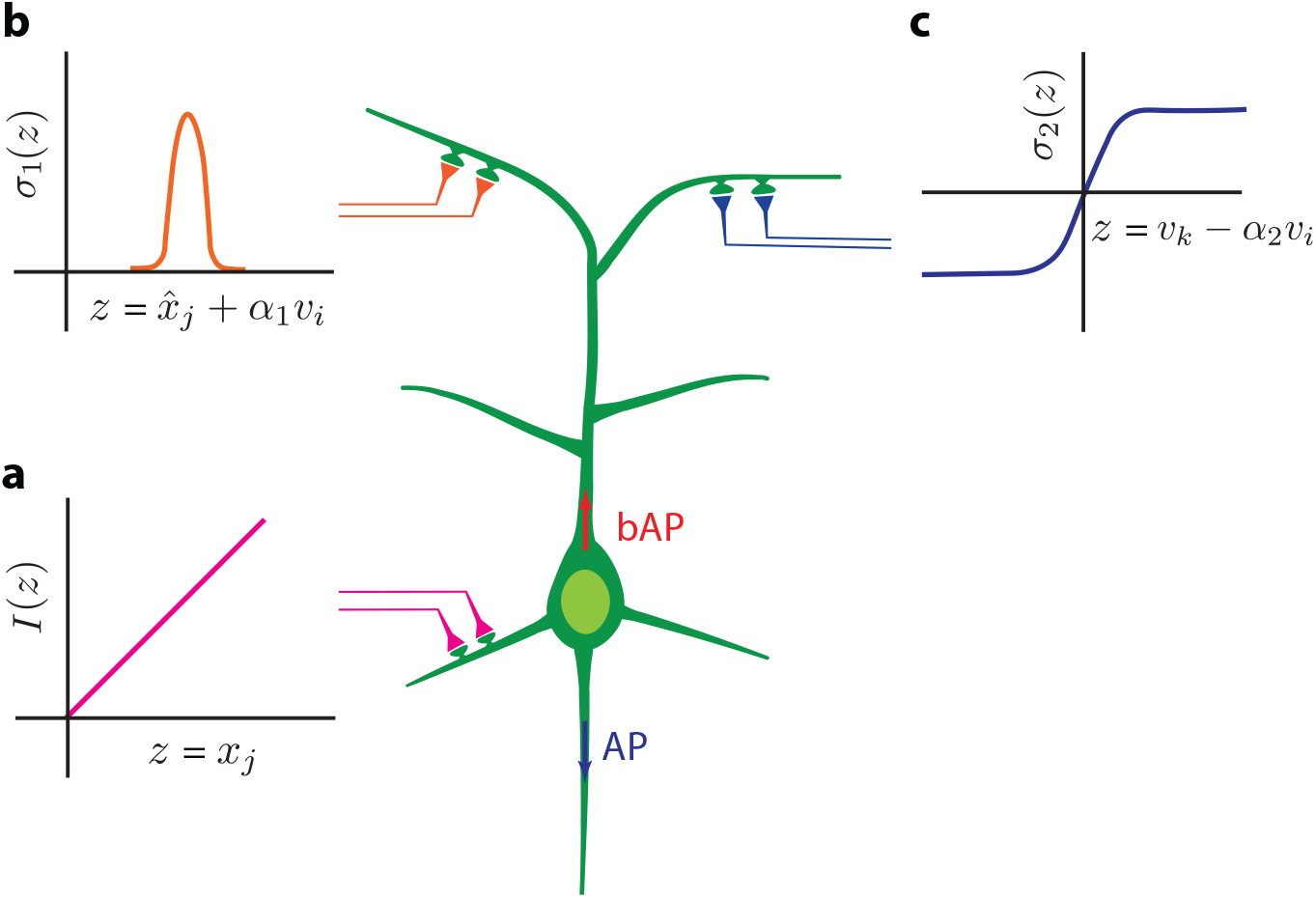
Schematic illustration of the proposed model. The proposed model (Eq. 1) can be considered as an extension of the point neuron model, where in addition to a term corresponding to linear integration of inputs (panel **a**), as in the standard model, two additional terms corresponding to different types of dendritic nonlinearities, both including bAPs, appear: dendrites with an XOR nonlinearity (panel **b**), and dendrites with a sigmoidal nonlinearity (panel **c**).

The second term in Eq. 1 represents the contribution of dendrites producing the XOR operation, whose existence has recently been reported (Gidon et al., 2020). When dendrites of this type receive inputs from two pathways, they produce a high response only when one of these pathways is active, i.e. the dendrites act as an XOR logical gate. In this term, the mean firing rate (*x*_*j*_) has been converted to the membrane potential 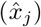, *α*_1_ is a scalar coefficient that represents the attenuation of the response (*v*_*i*_) backpropagating from the soma to the dendrite, and *σ*_1_ is a dendritic nonlinearity of XOR type, which for this reason must have a nonmonotonic form. We have represented this dendritic nonlinearity with a lobe (**Fig. 1b**): input values within the lobe indicate that just a single source is active, either 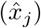 or (*α*_1_*v*_*i*_), and produce a positive response (XOR=1).

The third term in Eq. 1 includes lateral interactions between cortical units within a nonlinearity, *σ*_2_, that represents the contribution of sigmoidal-type dendrites (Jadi et al., 2014) (**Fig. 1c**). The inclusion of this term introduces a pool of different units that influence each other’s activity, similarly to other computational models emulating competition between neurons (Wilson and Cowan, 1972; Rozell et al., 2008). For unit *i* interacting with unit *k*, the input within the sigmoid nonlinearity includes the response of unit *k* (*v*_*k*_) which is subtracted from the backpropagating response of unit *i* (*v*_*i*_). As in the second term, *α*_2_ is a scalar that represents the attenuation of the backpropagating response (*v*_*i*_). The elements of *w*_2_ indicate the strength of the interactions between different units.

The mean firing rate of the neuron can be obtained by applying some nonlinear transform *φ* to the membrane potential *v*_*i*_ that is the solution of Eq. 1:

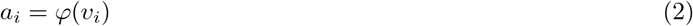

### Model instance

We define the scaling coefficients for each of the three terms in our model equation as follows:

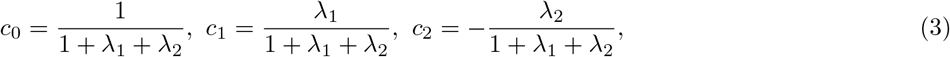

with *λ*_1_, *λ*_2_ ≥ 0.

In agreement with classical formulations (Field and Tolhurst, 1986), we select in the first term the linear filter *w*_0_ to be a Gabor function. For the second term, in line with experimental findings showing that dendrites with the XOR nonlinearity are apical (Gidon et al., 2020) and that the dendritic morphology in apical dendrites aligns with the orientation tuning of the neuron (Weiler et al., 2022), we select *w*_1_ to be a Gabor function with the same preferred orientation as *w*_0_, albeit with a different phase and Gaussian envelope. To convert firing rate (*x*_*j*_) into membrane potential 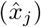 within the XOR nonlinearity, we perform full-wave rectification on *x*_*j*_ followed by exponentiation:

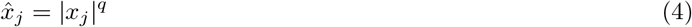

The elements of *w*_2_, in the third term, weigh the interactions between intracortical units. Again, following the classical view (Ben-Yishai et al., 1995; Rozell et al., 2008), we choose for *w*_2_ a function where *w*_2*ik*_ decreases as the difference between the preferred orientations of units *i* and *k* increases, essentially inducing the strongest competition between units with similar characteristics. The interaction weight of units *i* and *k* is equal to the absolute value of the inner product of their filters *w*_0_:

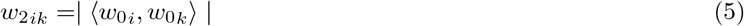

Finally, to convert the membrane potential output of our model to a spiking rate, we pass it through a rectified linear unit (ReLU), followed by exponentiation:

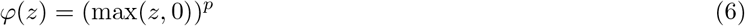

### Computing the model output

Finding the value *v*_*i*_ that solves Eq. 1 is not trivial, given that it appears on both sides of the equation: on the left side as the model output and on the right side as the bAP. To bypass this issue, we exploit the fact that solutions of Eq. 1 are steady states of the following differential equation:

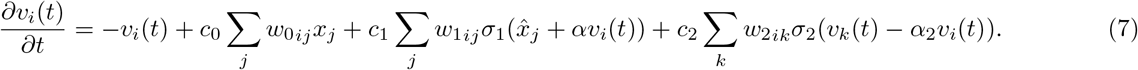

Although, in general, solutions to a differential equation do not necessarily converge to specific equilibria, this turns out to be true for Eq. 7, at least for *α*_2_ = 1. Moreover, the nonlinearities, *σ*_1_ and *σ*_2_, used in our model are locally integrable, which ensures the existence of their primitives (Fig. S1). Indeed, as we show in the supplementary material (Text S1), this equation is the gradient descent corresponding to an energy *E*(*v*), which has a unique minimum. This means that solutions of Eq. 7 are attracted to this minimum, which is then an equilibrium and hence the solution of Eq. 1. This suggests that the model is connected to the ecological modeling framework and is the result of a process aiming to optimize some particular feature, as we comment in the Discussion.

### Numerical implementation

The kernel of the linear term (*w*_0_) is a Gabor function positioned at the center of an *N*× *N* frame of gray background with *N* = 129 pixels. The grating’s period is one fourth the size of the frame (*L* = *N/*4 = 32.25 pixels) with an orientation coinciding with the unit’s preferred one (typically zero in our examples). The Gabor function is windowed by a Gaussian envelope with an aspect ratio (*γ*) equal to 0.125 and standard deviation (std = *L/*8 = 4) of 4 pixels. The kernel of the nonlinear XOR term (*w*_1_) is a Gabor function with the same orientation as *w*_0_, but phase-shifted by *π/*2 and a Gaussian window that is 3 times larger (std = 12 pixels). The kernel for the intracortical interaction term (*w*_2_) is computed from the linear term kernels of the different units (18 units with orientations spaced equally between 0 and 2*π*). The element *w*_2_(*j, k*), indicating the strength of the interaction between units *j* and *k*, is equal to the absolute value of the inner product of their linear kernels (Eq. 5).

The XOR nonlinearity is defined as follows:

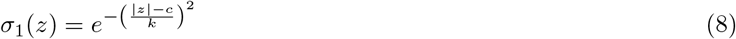

where *c* and *k* are constants that control the position of the XOR lobe and its half-width. Unless otherwise stated, *c* = 0.66 and *k* = 0.2.

The sigmoid nonlinearity, *σ*_2_, is defined as follows:

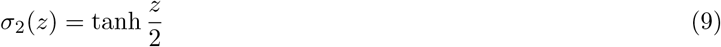

We numerically implement Eq. 7 via a forward Euler scheme with a time-discretization of Δ*t* = 0.1, and stop after a fixed number of iterations (*T* = 20). The initial condition is set at *v*_*i*_(0) = 0. An alternative stopping criterion based on estimating convergence (i.e. selecting *T* such that ∥*v*_*i*_(*T* + Δ*t*_*n*_) − *v*_*i*_(*T* )∥ *< ε* for some *ε >* 0) yields similar results.

### The spatial receptive field

Here, from the model equation (Eq. 1), we derive the spatial receptive field (RF) and show that it depends on the choice of stimulus, that is, it is input dependent, as indicated by experimental studies (Almasi et al., 2022; Butts, 2019). The RF is commonly defined in the literature as the input perturbation that produces a maximal output (Martinez-Garcia et al., 2018). From this definition, we derive a closed-form expression of the RF by taking the derivative of the model output with respect to the input (to simplify, we set *λ*_2_ = 0). For neuron *i*, its RF value at spatial location *j* is the following:

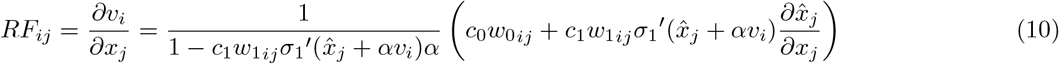

For *λ*_1_ = 0, the RF is equal to the filter *w*_0_, and as in the standard linear model, it does not depend on the input stimulus, while for *λ*_1_ ≠ 0, it becomes input-dependent due to the presence of the input, 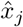, in the RF equation (Eq. 10). For instance, for two different stimuli, the model produces the same RF when its nonlinear term is zero (*λ*_1_ = 0) but two distinct RFs when there are nonlinear contributions (Fig. S2).

For *λ*_1_ *>* 0, the RF could receive, for certain inputs, contributions from outside of its classical RF part (*w*_0_; same as in the standard model), which could result in extra-classical RF effects. For example, we find that for *λ*_1_ ≠ 0, the model can reproduce the phenomenon known as endstopping (Hubel and Wiesel, 1968; Bolz and Gilbert, 1986; Rao and Ballard, 1999): the neural response to an optimally oriented input is diminished when the stimulus extends along the preferred orientation beyond a certain point (Fig. S3).

### Prolegomena to the experiments

Our model can explain a number of neurophysiological phenomena divided here into five experiments. Experiments 1-4 do not include intracortical interactions (*λ*_2_ = 0), but experiment 5 does. Box **1** provides summary information for each experiment: a short description, the model parameters investigated, and the inputs to the model. In the following, we also provide background information for experiment 3.

### Experiment 3: Volterra kernels, simpleness index, parametric fit, and input stimuli

From intracellular recordings in cat V1 neurons, Fournier et al. (2011) showed a dependence of simple- and complex-like functional components on visual stimulus statistics, with sparse noise stimulation steering neurons toward more complex-like receptive fields and dense noise directing them toward more simple-like ones. Stimulus-induced sub-threshold fluctuations and spiking responses were fitted in separate analyses with a nonparametric dynamic nonlinear model that includes first- and second-order Volterra kernels, the former representing the receptive field component that responds linearly to a stimulus contrast and the latter the component that is independent of contrast sign. Thus, the first-order Volterra kernel is considered the simple-like part of the receptive field, and the second-order one the complex-like. Simple and complex contributions for the different visual stimuli were quantified with the simpleness index (SI) measure, defined as the ratio of the spatiotemporal energy of the first Volterra kernel over the sum of the spatiotemporal energies of the first Volterra kernel and the diagonal entries of the second Voltera kernel. SI values toward zero indicate greater complex-like contributions, while SI values toward one indicate greater simple-like contributions, with a 0.5 value showing parity between the two.

We test the ability of our model to explain the experimental data of Fournier et al. (2011). We generate random inputs of sparse and dense noise of dimension 33 × 33. Sparse noise stimuli were constructed by randomly interspersing gray (same as the background), white, or black elements, with the average number of elements that contributed to the noise (white or black) being 7.26 out of the total 1089 elements. Dense noise stimuli were generated by randomly interspersing gray, white, or black elements with an equal probability of 0.33.

Analogously, we define SI for our simulations as the contribution of the first term of our model divided by its total response. The experimental SI data points (Fournier et al., 2011) followed the arc of a continuous smooth curve defined as follows:

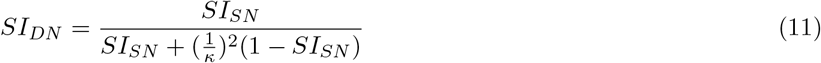

where *SI*_*DN*_ and *SI*_*SN*_ are the simpleness indices for dense and sparse noise, respectively, and *κ* is a free parameter. Using the Levenberg-Marquardt algorithm, we found the *κ* that provides the best fit to the data points produced by our model.

## Results

### Experiment 1: The model produces a continuum from simple to complex cell responses for thalamic input

To probe the capacity of the model to produce both simple and complex cell responses for thalamic input, we tested its response for different contributions of the XOR nonlinearity (*λ*_1_). For *λ*_1_ = 0, only the first linear term is nonzero and, consequently, the model’s response is modulated by the phase of the stimulus as one would expect from a typical simple cell (leftmost panel in **Fig. 2a**). As the nonlinear XOR contribution increases, the model becomes less sensitive to the phase of the stimulus, and eventually, for a sufficiently large *λ*_1_ value, it produces a phase-invariant response, showing the same orientation tuning irrespective of the phase of the stimulus (rightmost panel in **Fig. 2a**). The changes in the phase responses for 0 *< λ*_1_ *<* 10 indicate that by adjusting the contribution of the XOR nonlinearity the model can produce a continuum from simple to complex cell behavior, with further increases in *λ*_1_ having no noticeable effects (**Fig. 2b**).

**Figure 2:**
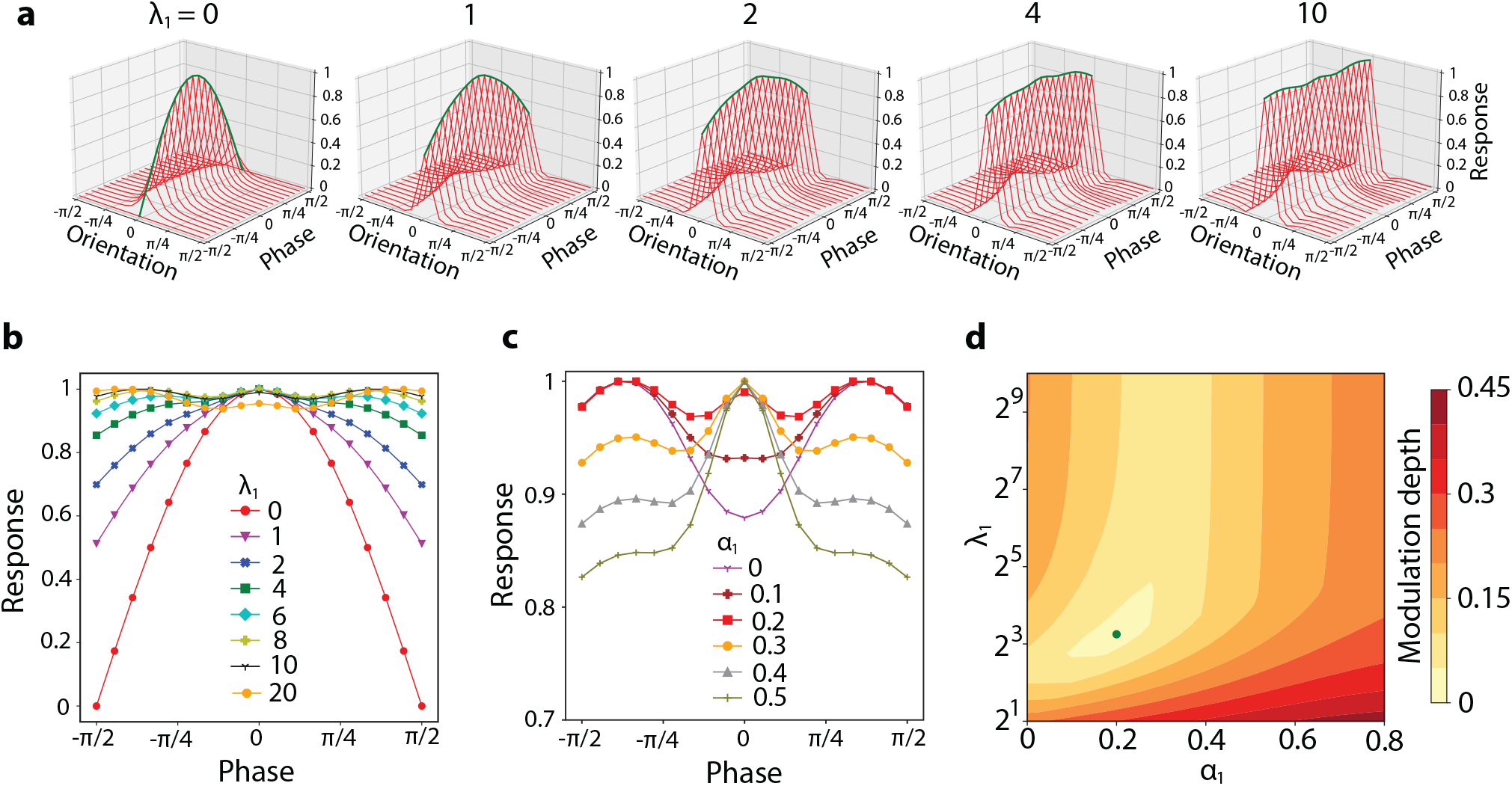
The XOR nonlinearity can produce a continuum from simple to complex cell behavior with the backpropagating response contributing to phase invariance. **a**: 3D plots of the model’s response as a function of stimulus orientation and phase for different *λ*_1_ values (*α*_1_ is fixed at 0.2). The green envelope shows the response of the model for stimuli oriented at 0 degrees (preferred orientation) at different phases (phase responses). **b**: Phase responses for different *λ*_1_ values (*α*_1_ is fixed at 0.2). **c**: Phase responses for different *α*_1_ values (*λ*_1_ is fixed at 10). **d**: Heatmap indicating the modulation depth value for different pairs of *λ*_1_ and *α*_1_ values.

In the analysis above, the weight of the backpropagating response was kept fixed (*α*_1_ = 0.2). To assess its effect in producing complex responses, we measured the model’s phase response for different values of *α*_1_. We found that the backpropagating response could have a constructive effect, since a nonzero contribution (*α*_1_ = 0.2) produces the flattest phase profile, with further increases having a detrimental effect (**Fig. 2c**). To quantify the degree of phase invariance for different combinations of *λ*_1_ and *α*_1_ values, we used the modulation depth metric, defined as the difference between the maximum and minimum responses in the phase response profile normalized by the maximum response. A complex cell with perfect phase invariance has a modulation depth of zero, with larger values indicating greater sensitivity to stimulus phase, i.e. simpler responses. We found that for *λ*_1_ = 10 and *α*_1_ = 0.2 the model produces the smallest modulation depth (**Fig. 2d**).

Our model provides an explanation for cells in V1 not being exclusively linear or nonlinear operators but instead producing intermediate behaviors; the greater the contribution of the XOR nonlinearity in a cell, the more phase-invariant (complex) its response is, with the backpropagating response contributing to its phase invariance. These results do not hold for arbitrary choices of the weight function *w*_1_. For instance, for a non-oriented *w*_1_ filter, the model loses its orientation selectivity (**Fig. 3a**). For a nonlinearity *σ*_1_ that is sigmoidal instead of type XOR, the model can still produce a complex response (**Fig. 3b**), but in this case the backpropagating response has no effect on its behavior (**Fig. 3c**).

**Figure 3:**
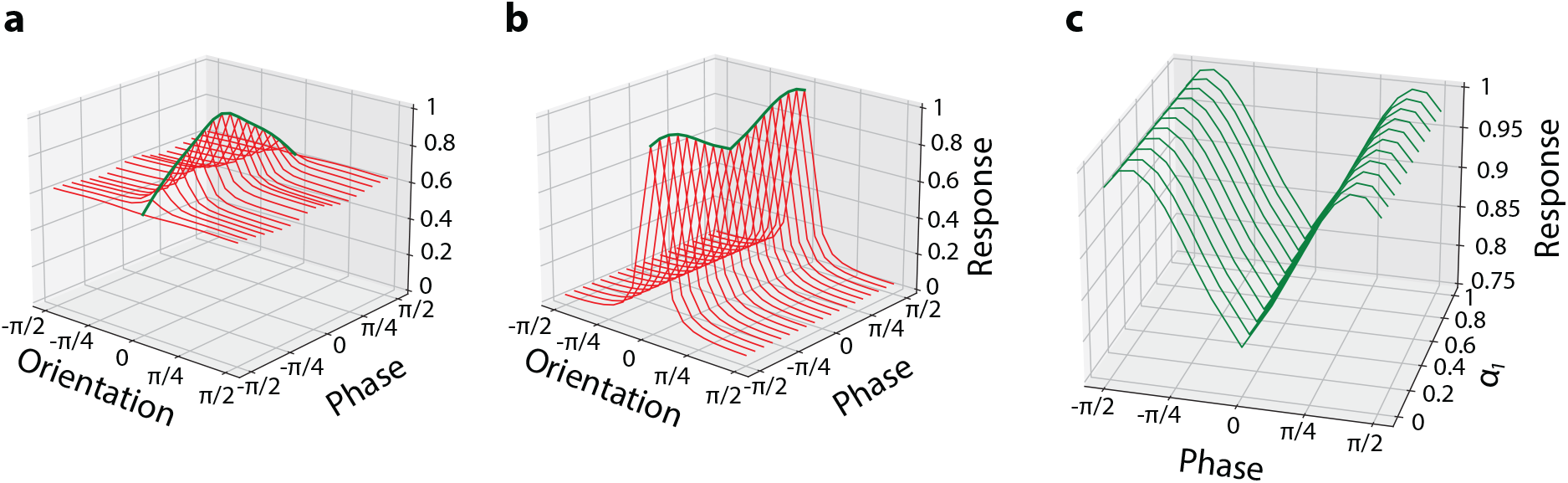
The model properties do not hold for arbitrary weight functions or nonlinearities. **a**: 3D plot of the model’s response as a function of stimulus orientation and phase for a non-oriented *w*_1_ filter (*λ*_1_ = 10 and *α*_1_ = 0.2). **b**: 3D plot of the model’s response as a function of stimulus orientation and phase for a nonlinearity *σ*_1_ that is sigmoid instead of XOR (*λ*_1_ = 10 and *α*_1_ = 0.2). **c**: 3D plot of the model’s phase responses for different *α*_1_ values for a sigmoid nonlinearity *σ*_1_.

These results show that the proposed model can explain complex cell responses for an input that is only thalamic, which is consistent with neurophysiology but a challenge for standard approaches. Of note, our model can also produce complex responses from simple cell inputs (Fig. S4), thus it is compatible as well with a hierarchical connectivity pattern.

### Experiment 2: The model is robust to perturbations

Nervous systems are robust to variability, as they can produce their intended function for a constellation of different configurations and parameter settings (Marom and Marder, 2023; Albantakis et al., 2024) and certain levels of noise inherent in their components (Faisal et al., 2008). In our model, noise-free Gabor filters, *w*_0_ and *w*_1_, suggest a specific alignment of the axonal branches of presynaptic neurons on the dendrites of the postsynaptic neuron. We tested the robustness of our model by adding different levels of noise to the filters, thus disturbing the alignment between neurites, and measuring its response. More specifically, to each filter we added a different instantiation of a noise matrix whose elements were random samples taken from a uniform distribution (the boundary values of the distribution are the minimum and maximum values of the filters). The noise matrices were also scaled by a constant representing the level of noise. We used parameter values for which the model produces a complex response (*λ*_1_ = 10 and *α*_1_ = 0.2). Qualitative inspection of the model’s response for different noise levels suggests that it is highly resilient, since even for added noise weights with high amplitude compared to the noiseless filter, the response pattern still appears complex (**Fig. 4a**). To confirm this observation, we compared the phase responses of the model with and without noise in their filters. We found that the mean square error between the two is close to zero even when the additive noise has an amplitude greater than twice the filters’ amplitude, with additional noise progressively degrading the response (**Fig. 4b**).

**Figure 4:**
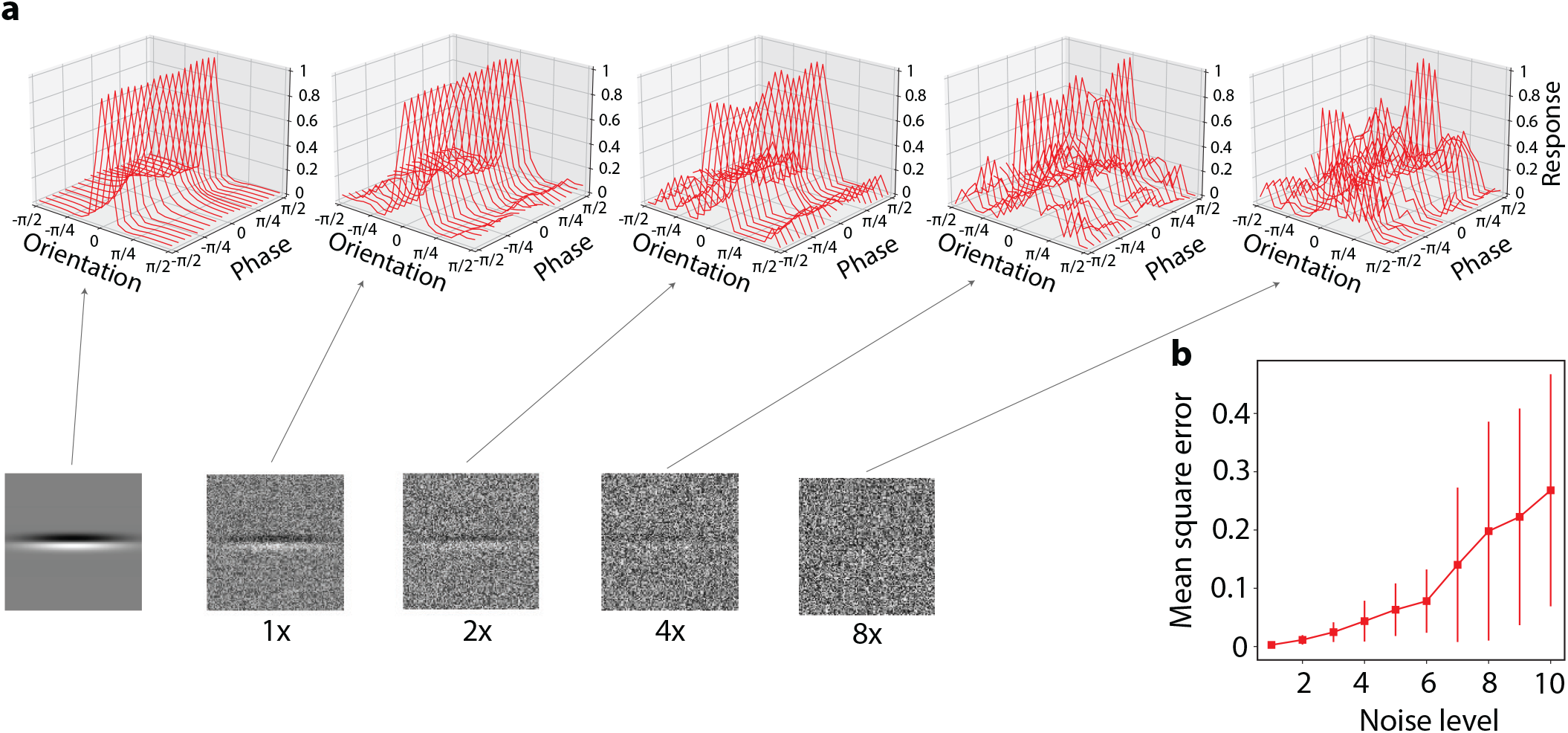
The model is robust to perturbations. **a**: 3D plot of the model’s response as a function of stimulus orientation and phase for different noise levels applied to filters *w*_0_ and *w*_1_. Examples of *w*_0_ filters with different noise levels are shown in the bottom row images. The arrows point to an example 3D plot for each noise level. Leftmost filter and 3D plot are with no noise. **b**: Mean square error between the phase responses of the model with and without noise, produced at each noise level. Each point and vertical line correspond to the mean and standard deviation from 100 repetitions. The correct phase response is considered the one produced by the model when the filters have no noise.

### Experiment 3: The model explains the transition of a neuron to simple or complex behavior with input statistics

From intracellular recordings in cat V1, Fournier et al. (2011) reported a puzzling phenomenon: the same cell has a more simple-like behavior for dense noise inputs, but a more complex-like one for sparse noise inputs. The authors fitted evoked neural responses with a second-order Volterra series expansion where the first- and second-order Volterra kernels represent the simple and complex part of the neurons’ receptive fields, respectively. They subsequently used a simpleness index (SI) metric to quantify the contribution of each part for sparse and dense noise stimulation: SI values toward zero indicate greater complex-like contributions, while SI values toward one indicate greater simple-like contributions (see Methods for a more detailed description). Bivariate analysis on the experimental SI data indicates that a neuron’s functional properties change with visual stimulation, with RF components appearing more simple-like in dense noise conditions than with sparse noise, as evidenced by the fact that all data points in **Fig. 5a** are above the identity line.

**Figure 5:**
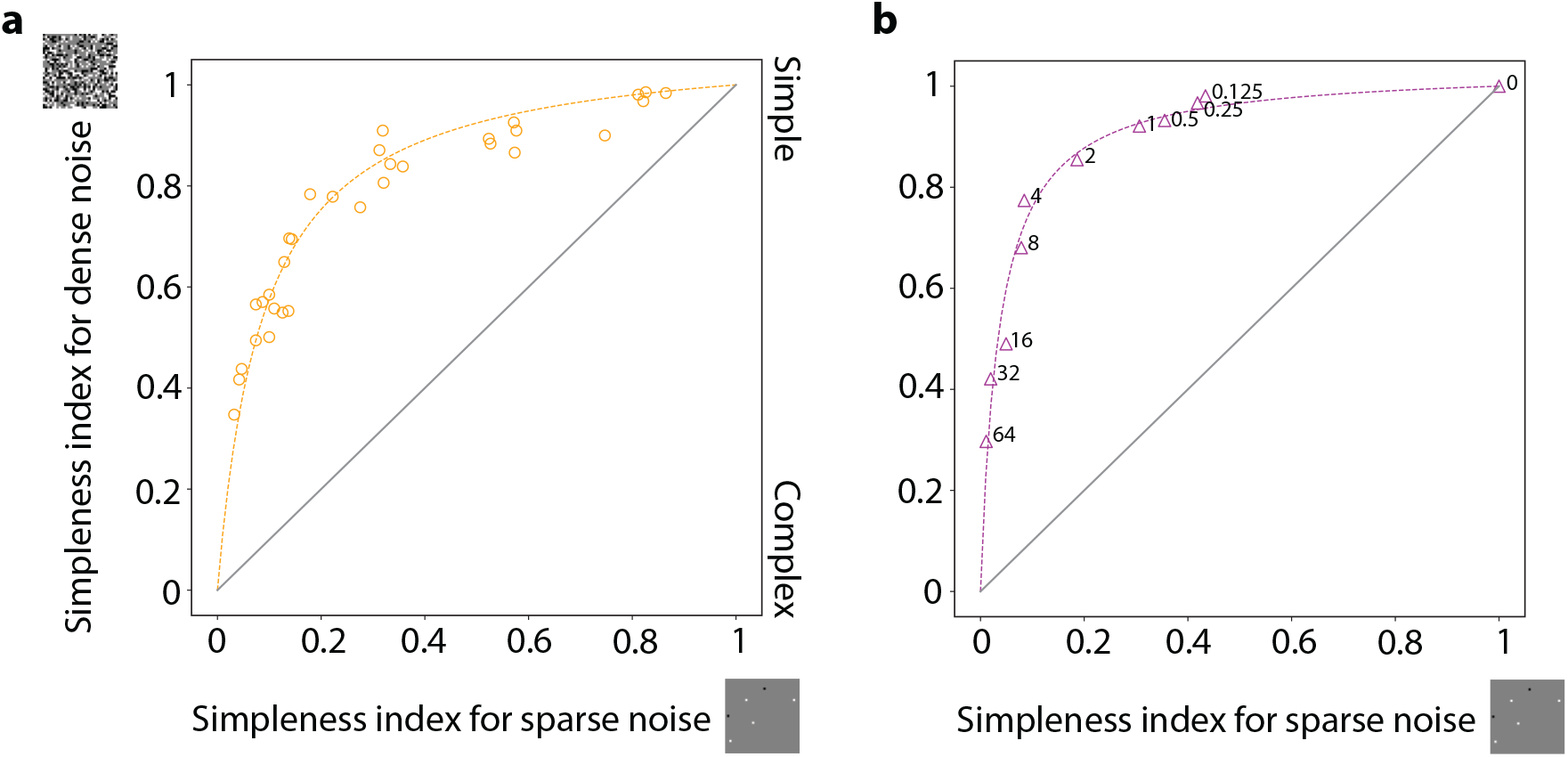
The model explains the transition of a neuron to simple or complex behavior with input statistics. **a**: Experimental data from fig. 3a in (Fournier et al., 2011). The plot shows SI for dense noise as a function of SI for sparse noise. Each open circle signifies the SI values for a single neuron estimated from subthreshold voltage fluctuations. The dashed curve is the best fit (*κ* value) to the data circles produced by Eq. 11. *κ* = 3.5 for the experimental data. **b**: Results for sparse and dense inputs for different instances of our model. We take *λ*_2_ = 0 and consider a wide range of values for *λ*_1_. We test each instance with 100 random inputs (for each noise condition) and compute the associated SIs, which are then averaged to produce the final SI value. The values of *λ*_1_ for each model instance are depicted next to their associated data point. The best fit (Eq. 11) for the data is for *κ* = 5.3.

To compare our model with neurophysiological data (Fournier et al., 2011), we consider its first two terms. We use a wide range of *λ*_1_ values, emulating for each parameter value the behavior of a neuron with different linear and nonlinear contributions. For all values of *λ*_1_, the model has a more simple-like behavior (larger SI value) for dense noise input than for sparse one (**Fig. 5b**). In general, our model has good qualitative correspondence with the experimental data, as evidenced by the positioning of the data points along the curve produced by Eq. 11 (compare **Fig. 5a** with **Fig. 5b**).

Fournier et al. (2011) explained this behavior with a model that changes its parameters with the input stimulus, and conjectured that sparse stimuli recruit lateral interactions that contribute to complex-like nonlinearities. Our model, on the other hand, does not change for different inputs and can reproduce the effect solely with thalamic input (fixed parameters and *λ*_2_ = 0), relying on the XOR nonlinearity. For no contribution from XOR dendrites (*λ*_1_ = 0), SI is 1 for both types of input, indicating simple cell behavior, and as *λ*_1_ increases, the cell tends towards a more complex-like behavior, in agreement with what was observed in **Fig. 2**.

### Experiment 4: The model explains the relationship between phase sensitivity and input contrast for complex cells

Recordings in V1 show that complex cell responses become less phase invariant as stimulus contrast decreases (Crowder et al., 2007; Van Kleef et al., 2010). This property is more common in superficial cortical layers (Meffin et al., 2015), which receive most of the thalamic input, but is not as prominent in deeper layers, where the input is mainly from cortical (and likely orientation selective) cells. To probe how our model compares with experimental data, we tested its degree of phase sensitivity for different stimulus contrasts using the modulation depth metric. For thalamic input, our model can achieve good qualitative correspondence with respect to experimental data with a fixed set of parameters and for a wide range of *λ*_1_ values, since for both the model and the experimental data (Meffin et al., 2015), modulation depth decreases linearly as stimulus contrast increases logarithmically (**Fig. 6a**). To obtain a fit with the experimental data, we adjusted the exponents that convert the firing rate to membrane potential inside the nonlinearity in the second term (from *q* = 0.5 in all other experiments to *q* = 0.25 here; Eq. 4), and the membrane potential output of the model to spiking rate (from *p* = 1 in all other experiments to *p* = 0.5 here; Eq. 6). Experimental evidence indicates that our choices are within the range of acceptable values (Mechler and Ringach, 2002). The model’s modulation depth still decreases linearly, albeit for a smaller range of contrasts, for no adjustments on the exponents (Fig. S5).

**Figure 6:**
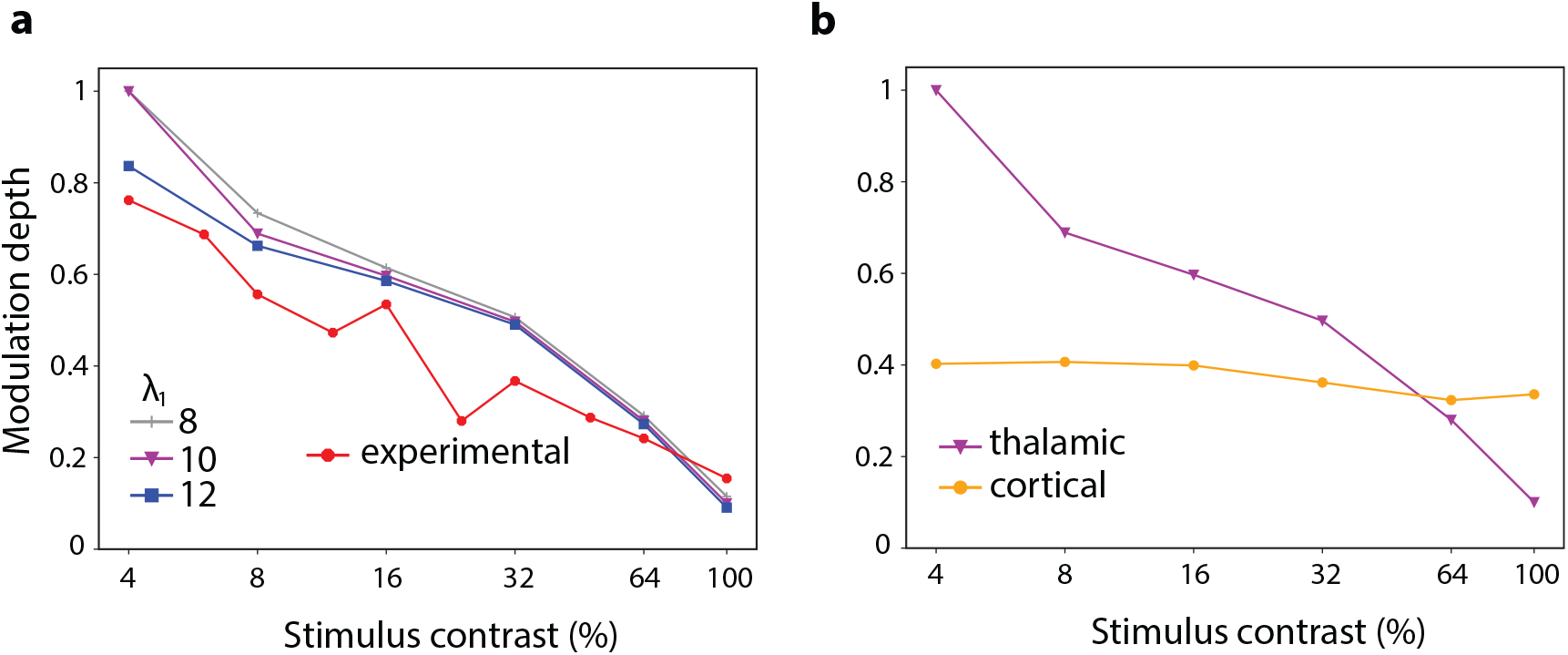
The model, with fixed parameters, explains the relationship between stimulus contrast and phase sensitivity shown experimentally. **a**: Modulation depth as a function of stimulus contrast, for different values of *λ*_1_ overlaid with experimental data taken from fig. 3c in (Meffin et al., 2015). **b**: Same as in **a** for two different instances of our model. In both cases *λ*_1_ = 10, however for the thalamic condition the input *x* is as in **a** while for the cortical condition *x* is filtered with a Gabor function with the same orientation preference as *w*_0_.

The dependence of a V1 neuron’s phase sensitivity on stimulus contrast is not as prominent for mice compared to primates and cats (Yunzab et al., 2019b) and virtually disappears for V2 neurons in macaques (Cloherty and Ibbotson, 2015). In both cases, the input passes through oriented filters: V1 cells in mice receive thalamic input that is orientation selective (Zhao et al., 2013), and V2 cells in macaques receive the majority of their input from V1 (Felleman and Van Essen, 1991; Markov et al., 2014). To test whether our model can produce this behavior, we mimicked the effect of orientation-selective thalamic or V1 cells by passing the input *x* through an oriented filter. As in the physiological results, the phase sensitivity of our model this time did not depend on stimulus contrast (**Fig. 6b**). In the hierarchical case, we set *q* = 0.2, and also adjusted the position of the lobe in the XOR nonlinearity, with minor adjustments from our preset values, also producing a similar trend.

### Experiment 5: Intracortical interactions sharpen orientation tuning

To account for the prominent local intracortical connections within V1, we introduce a third term in the model (*λ*_2_ ≠ 0), different from the ones used in other well-known computational models (Wilson and Cowan, 1972; Hopfield, 1984; Rozell et al., 2008), in that it includes a backpropagating response. The nonlinearity we use for the cortical interaction term is the sigmoid function, a choice that is consistent with the response profile of a subset of dendrites (Jadi et al., 2014). In line with a number of computational studies (Somers et al., 1995; Ben-Yishai et al., 1995; Troyer et al., 1998; McLaughlin et al., 2000; Wielaard et al., 2001; Shelley et al., 2002; Shapley et al., 2003), we hypothesized that lateral cortical interactions modulate orientation tuning.

We first tested whether the inclusion of the cortical interaction term interferes with the ability of the XOR non-linearity to produce a continuum from simple to complex cell responses. To that end, we kept the cortical interaction term fixed (*λ*_2_ = 1) and varied the contribution of the XOR nonlinearity (*λ*_1_). We found that the cortical interaction term does not disrupt the action of the XOR nonlinearity, since increasing the value of *λ*_1_ produces a gradual change from simple to complex cell responses (**Fig. 7a**), akin to what was shown for *λ*_2_ = 0 (**Fig. 2a**).

**Figure 7:**
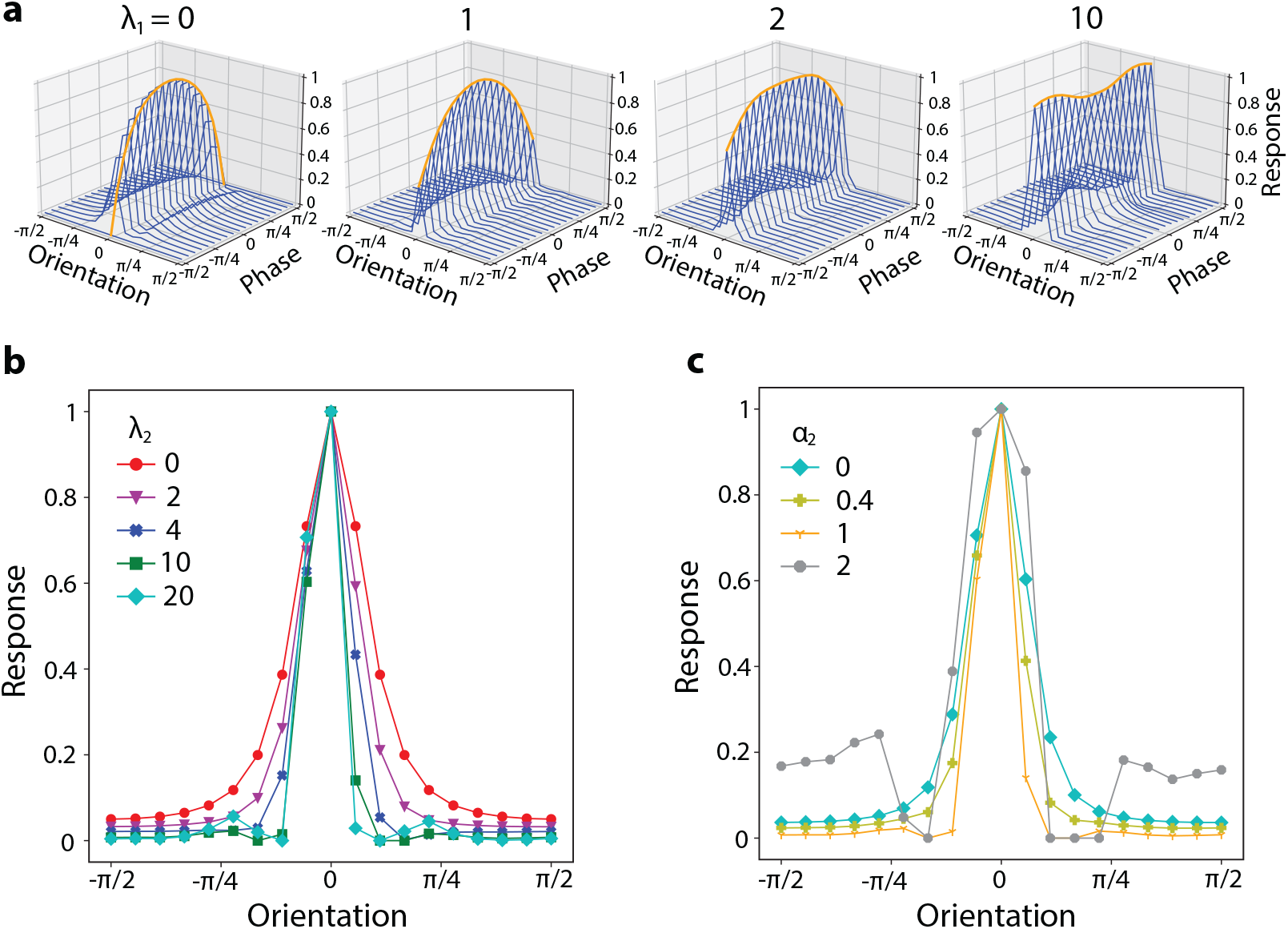
Cortical interactions in the model sharpen orientation tuning curves with the backpropagating response contributing to it. **a**: 3D plot of the model’s response as a function of stimulus orientation and phase for a fixed contribution from the cortical interaction term (*λ*_2_ = 1) and variable contributions of the XOR nonlinearity (*λ*_1_). Our tests consider 18 neural units, with orientation selectivities equally spaced between 0 and 2*π* that interact through the action of the cortical interaction term. **b**: Model’s response as a function of stimulus orientation (for a fixed optimal phase) as we vary the strength of the cortical interaction term (*λ*_2_), with the rest of the parameters being fixed (*λ*_1_ = 10, *α*_1_ = 0.2, *α*_2_ = 1). **c**: Model’s response as a function of stimulus orientation (for a fixed optimal phase) as we vary the strength of the backpropagating response in the cortical interaction term (*α*_2_), with the rest of the parameters being fixed (*λ*_1_ = 10, *α*_1_ = 0.2, *λ*_2_ = 10).

For a fixed XOR contribution (*λ*_1_ = 10), we observe that increases in the strength of the cortical interaction term within a certain range (0 *< λ*_2_ *<* 10) progressively induce a sharpening of the orientation tuning curve (**Fig. 7b**). For this analysis, the contribution of the backpropagating response within the intracortical term was fixed (*α*_2_ = 1). We subsequently kept the cortical interaction term fixed (*λ*_2_ = 10) and varied the strength of its backpropagating response (*α*_2_) to test the latter’s effect. We found that the backpropagating response has a constructive effect in the sharpening of orientation tuning up to a certain value, with further increases degrading the orientation tuning curve, for instance, for *α*_2_ = 2 (**Fig. 7c**).

## Discussion

We have proposed a model for V1 that considers nonlinear dendritic integration and bAPs. The model can be viewed as an extension of the standard model, minimizes an energy function, allows for a better conceptual understanding of neural processes, and explains several neurophysiological phenomena that have challenged classical approaches.

### A variational dynamical model

The dynamical formulation in Eq. 7 allows us to connect our model to the ecological modeling framework, which formulates efficient coding as an optimization problem. The solutions are then obtained using evolution equations that minimize an energy functional (Atick, 1992). In the field of mathematical optimization, this is known as a variational formulation, and the resulting solutions achieve optimality through minimal redundancy.

The existence of an energy underlying the dynamical formulation of Eq. 7 emerges naturally from the proposed equation. This is a desirable feature because while a variational principle invariably produces an evolution equation, the converse does not hold: not all evolution equations correspond to the minimization of an energy functional. This is the case, for example, for neural field models and other dynamical frameworks that describe the processing of stimuli in the visual cortex (Wilson and Cowan, 1972; Chow and Karimipanah, 2020), which do not satisfy any variational principle (Bertalmío et al., 2021), thereby being suboptimal for redundancy reduction. In that sense, Eq. 7 can be seen as a generalization of the classical Wilson-Cowan (Chow and Karimipanah, 2020) or the more recent ORGaNICs formulation (Heeger and Mackey, 2019), now with different types of nonlinearities that are input dependent; our model is also closely related to the LHE and INRF methods (Bertalmío et al., 2020, 2021). Moreover, the variational nature of the model allows for improved interpretability.

The energy functional minimized by Eq. 7 is the sum of three terms, each accounting for different minimization criteria. The first term is an orientation-dependent term, present in standard linear filtering models. The second term, the main novelty of the model, is both input- and orientation-dependent and accounts for the simple to complex cell behavior. The last term is input-independent, favoring variability in the neural response, establishing, with its orientation tuning effect, a direct connection to the sparse coding framework, a family of energy-inspired vision models (Olshausen and Field, 1996; Rozell et al., 2008; Paiton et al., 2020; Rentzeperis et al., 2023; Oldenburg et al., 2024).

### The XOR term and other dendritic nonlinearities

A recent study on pyramidal neurons in the human cortex has revealed a new class of calcium-mediated dendritic channels that could solve the XOR problem (Gidon et al., 2020), with a subsequent study showing evidence of dendrites with similar properties in rats (Magó et al., 2021). What makes these dendritic channels distinct from any other studied is that they produce a high response when one of two groups of synaptic inputs is active, and a suppressed response when both or none are. Thus, uniquely, this dendritic nonlinearity is represented with a non-monotonic function, and by varying its contribution our model can produce a continuum from simple to complex cell responses (**Fig. 2a-b**).

Due to their varied biophysical properties, dendrites offer a large repertoire of input transformations. We have opted to use the XOR nonlinearity for our feedforward input, though we must note that the more commonly described nonlinearities in pyramidal cell dendrites are both monotonic and saturating –for example, nonlinearities that are mediated by sodium ion channels (Stuart et al., 1997; Golding and Spruston, 1998; Losonczy and Magee, 2006), or that are N-Methyl-D-aspartate (NMDA) receptor-dependent (Antic et al., 2010; Major et al., 2013). The flexibility of our model allows us to use these nonlinearities as well; in fact, in this study, we replaced the XOR nonlinearity with a sigmoidal one, and the model could still reproduce complex responses (**Fig. 3b**). By using other monotonic functions or variations of the sigmoid, our model could probe in a systematic way the more well-established dendritic types.

### The role of backpropagating action potentials

Action potentials (APs) initiated at the axonal initial segment travel down the axon of a neuron, but can also propagate back to its dendrites (Stuart and Sakmann, 1994; Stuart et al., 1997). Due to their unclear role, bAPs may appear as an offshoot of APs. Their interaction with the dendritic membrane potential can vary, promoting either an increase in subsequent spiking generation (Larkum et al., 1999) or a decrease (Golding and Spruston, 1998; Remy et al., 2009), with possible ramifications in plasticity and learning (Feldman, 2012).

In our formulation, backpropagating responses are included in the XOR nonlinearity that receives feedforward input and the sigmoid nonlinearity that processes the interacting intracortical units. In both cases, the backpropagating response has a constructive effect; it promotes phase invariance in the XOR term (**Fig. 2c-d**), and it sharpens the units’ orientation tuning curve in the intracortical term (**Fig. 7c**).

Our aim in this study was to construct a general model that could capture some principal aggregate effects of dendritic processing. Under this reductionist formulation, the model can represent some of the underlying neural dynamics since it can explain a number of phenomena that remain challenging for current models under the standard framework and also produce working hypotheses for experimental studies. For an examination of its putative role on plasticity, a simplified formulation of bAPs, similar to ours, can be incorporated in adaptive rewiring models of the brain (Rubinov et al., 2009; Rentzeperis et al., 2022; Lynn et al., 2024).

### The role of connectivity

Anatomical studies show that a minority of the input to V1 cells is from geniculate cells (feedforward), while a significant portion of cortical connections is lateral and recurrent (Binzegger et al., 2004; Peters et al., 1994; Gilbert and Wiesel, 1983).Here, we incorporate a term that describes recurrent interactions between units and results in sharpening of their orientation tuning curves (**Fig. 7b**). Different models of V1 processing have also employed lateral cortical interactions along with feedforward input to explain orientation selectivity (Somers et al., 1995; Ben-Yishai et al., 1995; Troyer et al., 1998; McLaughlin et al., 2000; Wielaard et al., 2001; Shelley et al., 2002; Shapley et al., 2003)-with another study, however, contesting this functionality (Seriès et al., 2004). Due to its modular nature, our model can test a host of functions mediated by different connectivities as proposed by experiments (Olsen et al., 2012; Gilbert and Li, 2013; Harris et al., 2019; Keller et al., 2020).

### Conclusion and future work

We have developed a formulation that can be seen as a contribution in the search for a consensus model for V1 cell responses (Martinez and Alonso, 2003). It includes general dendritic mechanisms and functionalities from different processing hierarchies to account for a neuron’s nonlinear behavior and its anatomical connections. Our model offers several conceptual advantages. First, a better understanding of neural processes through a simple, yet flexible, formulation that can explain various phenomena with a reduced and fixed set of parameters. Second, it is modular and adaptable: new terms representing other types of dendrites and functional connectivity patterns can be added, and the elements of each term can be updated as new experimental findings become available. Third, it could help our understanding of the role of bAPs in neural processing.

Most current modeling approaches are data-driven, with an emphasis on improving predictions, often at the expense of biological interpretability (Frégnac and Bathellier, 2015). In recent years, the call for better conceptual frameworks has increased (Olshausen, 2013; Frégnac, 2017; Saxe et al., 2021), as models whose mathematical form is based on physiological mechanisms can disentangle complex interactions, revealing localized elements within the model that would otherwise be distributed over several subunits in current approaches (Butts, 2019). Our model strives in that direction.

While our experiments have explored the case of cortical cells receiving an input that is either thalamic or cortical, the weight of the evidence suggests that thalamic input is, numerically, only a small fraction of synaptic input onto V1 pyramidal cells. Future work can test our model in the scenario where the inputs to the first nonlinear term come from both LGN and V1, and the dendritic nonlinearity is of a different type. The model can also be extended so that it can handle time-dependent stimuli, e.g. by treating as physical time the evolution variable in Eq. 7, as is often done in neural field models (Wilson and Cowan, 1972; Chow and Karimipanah, 2020) or by introducing temporal filters in the model, as was recently done for the closely related INRF model (Luna et al., 2025). The model could potentially be useful for machine learning applications in which the model weights could be learned by training an artificial neural network, following the approach that Rawat et al. (2024) took for the dynamical neural framework in (Heeger and Mackey, 2019).

## Code availability

The source code with the implementation of the model and the generation of all figures is available on GitHub: https://github.com/rentzi/nonlinearNeuronModel

## Acknowledgements

We thank Luca Calatroni and Valentina Franceschi for very inspiring discussions that helped in the initiation of this project, and Steeve Laquitaine and Panayiota Poirazi for comments on the manuscript. For this project, we acknowledge the funding of PID2021-127373NB-I00, Ministry of Science and Innovation (Spain); VIS4NN - Programa Fundamentos Fundación BBVA 2022 (Spain); IDEALCV-CM (ref. TEC 2024/COM-322), Comunidad de Madrid (Spain); ANR-20-CE48-0003 and ANR-11-IDEX-0003-02, National Research Agency (France).

## Author contributions

IR: data analysis, methodology, investigation, and writing. DP: methodology, investigation, and writing. MB: conceptualization, data analysis, methodology, investigation, and writing.

## List of Tables

**Table 1:**
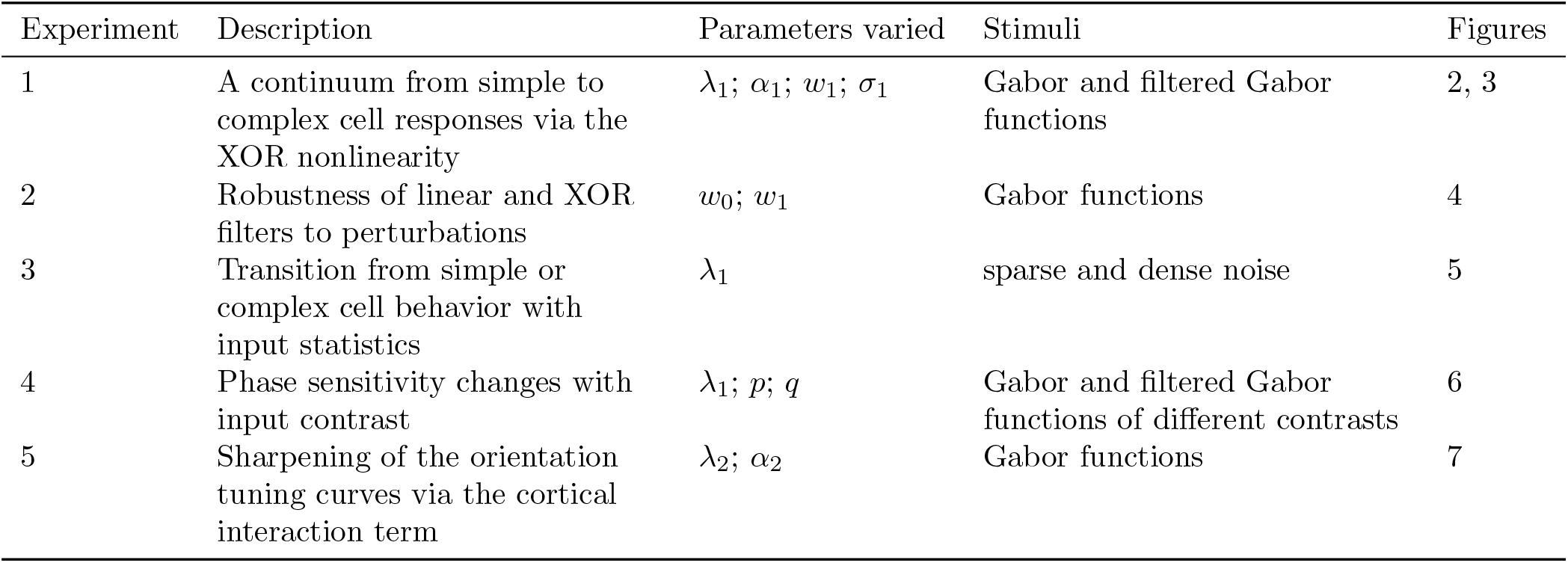
Overview of the experiments. Each row provides information for an experiment. The first column gives the experiment number; the second, a brief description of the physiological phenomenon the model explains; the third, the parameters of the model that were probed. Note that from Experiments 1 to 4 the cortical interaction term is not included (*λ*_2_ = 0); Experiment 5 includes it and varies its contribution to probe its effect in the response of the model. The fourth column indicates the input used. For brevity’s sake, when we write Gabor functions we mean a set of Gabor stimuli of the same frequency but different preferred orientations and phases. Filtered Gabor functions refer to the same set of stimuli but convolved with another Gabor function to simulate the first step of the hierarchical model; i.e. the input passing through a simple V1 cell. The last column refers to the Figures each experiment corresponds to.

| Experiment | Description | Parameters varied | Stimuli | Figures |
| --- | --- | --- | --- | --- |
| 1 | A continuum from simple to complex cell responses via the XOR nonlinearity | $\lambda_1; \alpha_1; w_1; \sigma_1$ | Gabor and filtered Gabor functions | 2, 3 |
| 2 | Robustness of linear and XOR filters to perturbations | $w_0; w_1$ | Gabor functions | 4 |
| 3 | Transition from simple or complex cell behavior with input statistics | $\lambda_1$ | sparse and dense noise | 5 |
| 4 | Phase sensitivity changes with input contrast | $\lambda_1; p; q$ | Gabor and filtered Gabor functions of different contrasts | 6 |
| 5 | Sharpening of the orientation tuning curves via the cortical interaction term | $\lambda_2; \alpha_2$ | Gabor functions | 7 |

