## Supplementary material for "A neural model for V1 that incorporates dendritic nonlinearities and back-propagating action potentials"

#### Text S1

##### Energy formulation

We rewrite the differential equation used to compute the model (Eq. 7 in the main text) as

$$\dot{v} = \frac{\partial v}{\partial t} = F_0(v) + c_1 F_1(v) + c_2 F_2(v), \tag{1}$$

where the vector fields $F_{\{0,1,2\}}$ are defined by

$$F_0(v)_i = -v_i + c_0 \sum_j w_{0ij} x_j, \quad F_1(v)_i = \sum_j w_{1ij} \sigma_1(\hat{x}_j + \alpha v_i), \quad F_2(v)_i = \sum_j w_{2ij} \sigma_2(v_j - v_i), \tag{2}$$

and we have set $\alpha_2 = 1$ so now we call $\alpha_1$ simply $\alpha$. In this section, we consider the following minimal assumptions on the parameters: we have that $w_{2ij} = w_{2ji}$, and that $\sigma_2$ is an odd function. We also assume that $\sigma_1$ and $\sigma_2$ are locally integrable, which ensures the existence of their primitives $\Sigma_1$ and $\Sigma_2$, respectively:

$$\Sigma_i(s) = \int_0^s \sigma_i(r)\, dr, \qquad i = 1, 2.$$

Observe that $\Sigma_1(0) = \Sigma_2(0) = 0$. Although the following is independent of the exact form of $\sigma_1$ and $\sigma_2$, we present the graphs of the instances that we chose for the experiments, as well as their corresponding primitives, in Figure S1.

The aim of this section is to connect Eq. 1 with the gradient descent corresponding to the energy

$$E(v) = E_0(v) - c_1 E_1(v) - c_2 E_2(v), \tag{3}$$

where

$$E_0(v) = \frac{1}{2} \sum_i \left| v_i - c_0 \sum_j w_{0ij} x_j \right|^2, \quad E_1(v) = \frac{1}{\alpha} \sum_{ij} w_{1ij} \Sigma_1(\hat{x}_j + \alpha v_i), \quad E_2(v) = \frac{1}{2} \sum_{ij} w_{2ij} \Sigma_2(v_i - v_j). \tag{4}$$

Observe that each of the three terms above accounts for different minimization criteria to which the neural coding is subjected. The first term, $E_0$, is quite standard and its minimizer is $v_i = c_0 \sum_j w_{0ij} x_j$. That is, this energy induces a linear filtering of the input.

The second term $E_1(v)$ in the definition of $E(v)$ accounts for the first non-linear term appearing in Eq. 1. Due to the negative sign, it must be maximized. Its maximum can be explicitly computed and is given by

$$v_i = \arg\max_u \sum_j w_{1ij} \Sigma_1(\hat{x}_j + \alpha u). \tag{5}$$

That is, this energy selects $v_i$ as a bias $u$ for the function $\Sigma_1$ (which can be thought of as being a piece-wise non-decreasing function) in such a way that the response of the oriented Gabor filter $w_1$ to the stimulus $\Sigma_1(\hat{x}_j + \alpha u)$ is maximal. As the experiments show, this is the term that accounts for the simple to complex cell behavior.

Finally, the third term $E_2(v)$, which must be maximized, is input-independent and favours the variability of the neural response. In fact, since $w_{2ij}$ is positive and decreases with the difference in the preferred orientation at $i$ and $j$, it follows from the shape of $\Sigma_2$ that maximizing this energy corresponds to choosing $v_i$ and $v_j$ as different as possible.

We then have the following.

**Proposition 1.** *Eq. 1 is the gradient descent equation corresponding to the energy $E(v)$. That is, for any neuron $v_i$ and any time $t$ it holds*

$$\dot{v}_i(t) = -\frac{\partial E(v(t))}{\partial v_i}. \tag{6}$$

*Moreover, the energy $E$ admits a unique minimum.*

*Proof.* Since the derivative is a linear operator, it suffices to show that $-\partial E_r/\partial v = F_r(v)$ for $r = 0, 1, 2$. The cases $r = 0$ and $r = 1$ follow directly by differentiating the expressions for $E_0$ and $E_1$. On the other hand, for any index $k$, by the oddness of $\sigma_2$, and the fact that $w_{2ik} = w_{2ki}$, we have that

$$\begin{aligned} -\frac{\partial E_2(v)}{\partial v_k} &= \frac{1}{2}\left[\sum_j w_{2kj}\hat{\sigma}_2(v_k - v_j) - \sum_i w_{2ik}\hat{\sigma}_2(v_i - v_k)\right] \\ &= \sum_j w_{2kj}\hat{\sigma}_2(v_k - v_j) \\ &= F_2(v)_k. \end{aligned} \tag{7}$$

To complete the proof of the statement, we are left to show the existence of a global minimum. Since $v$ belongs to the finite dimensional space $\mathbb{R}^N$, this follows from a lower bound of the form

$$E(v) \geq \|v\|_2^2 - a\|v\|_2 - b, \tag{8}$$

for some $a, b \in \mathbb{R}$. Here, we let $\|v\|_2^2 = \sum_i |v_i|^2$. Indeed, Eq. 8 implies that $E(v) \to +\infty$ as $\|v\|_2 \to +\infty$, and hence there exists $M > 0$ such that the infimum of $E$ on $K = \{\|v\|_2 \leq M\}$ is the same as the one on the full space $\mathbb{R}^N$. The continuity of $E$ and the compactness of $K$ then yield the statement.

Developing the square in the definition of $E_0$, and letting $z_i = c_0 \sum_j w_{0ij} x_j$ we have

$$2E_0(v) = \sum_i |v_i|^2 + 2\sum_i z_i v_i + \sum_i |z_i|^2 \geq \|v\|_2^2 - a_0\|v\|_2 - b_0.$$

Here, we used Cauchy-Schwarz inequality and let $a_0 = 2\|z\|_2$ and $b_0 = \|z\|_2^2$.

On the other hand, since $\Sigma_1(s) \leq \|\sigma_1\|_\infty |s|$, there exists $a_1, b_1 > 0$ such that

$$|E_1(v)| \leq \frac{\|\sigma_1\|_\infty}{|\alpha|}\|w_1\|_\infty N\left[\sum_j |\hat{x}_j| + |\alpha|\sum_i |v_i|\right] \leq a_1 + b_1\|v\|_2.$$

Here we used the fact that $\sum_i |v_i| \leq \sqrt{N}\|v\|_2$ for any $v \in \mathbb{R}^N$. Finally, a similar argument exploiting the fact that $\Sigma_2(s) \leq \|\sigma_2\||s|$ yields that there exists $a_2, b_2 > 0$ such that

$$|E_2(v)| \leq a_2 + b_2\|v\|_2.$$

The lower bound in Eq. 8 then follows by putting together the above estimates and letting $a = \max\{a_0, a_1, a_2\}$ and $b = \max\{b_0, b_1, b_2\}$. □

#### Supplementary Figures

### List of Figures

S1 Graphs of the non-linearities $\sigma_1$ and $\sigma_2$ used in the experiments, together with the corresponding primitives $\Sigma_1$ and $\Sigma_2$ . . . 5

S2 **The model has the same RF when its nonlinear terms are zero ($\lambda_1 = \lambda_2 = 0$) irrespective of the input stimulus, but for nonzero nonlinear contributions, its RF becomes stimulus dependent.** Results for a horizontal (top row) and a vertical (bottom row) sinusoidal grating stimulus (in all cases, $\lambda_2 = 0$). First column shows the stimuli, the other columns, the RFs for different $\lambda_1$ values. For $\lambda_1 = 0$, the RF does not depend on the stimulus and is the same in both cases, for $\lambda_1 \neq 0$ it is stimulus dependent. . . . 6

S3 **For nonzero nonlinear contributions ($\lambda_1 = 1$), the model shows endstopping.** Normalized spiking response of the model (omitting the second nonlinear term, i.e. with $\lambda_2 = 0$) for three stimuli: $S1$ (part of a Gabor presented only on the sides, outside of the classical RF), $S2$ (a Gabor presented at the center, inside the classical RF), and their sum, $S1 + S2$. The red diamond markers indicate the response of the unit when $\lambda_1 = 0$, and the green circle markers when $\lambda_1 = 1$. The images above the markers indicate the RF of the unit for the $\lambda_1$ value corresponding to the marker below and the stimulus corresponding to the abscissa position. For no nonlinear contribution ($\lambda_1 = 0$), the RF stays the same irrespective of the stimulus and the model does not show endstopping; the responses for $S2$ and $S1 + S2$ are the same. For a contribution from the nonlinear term ($\lambda_1 = 1$), the RF changes with the stimulus, and the response of the unit shows endstopping; it decreases for the elongated stimulus ($S1 + S2$) compared to the Gabor within the classical RF ($S2$), but does not respond to the side parts of the stimulus ($S1$) when they are presented in isolation. . . . 7

S4 **The model produces a complex response with cortical input.** 3D plot of the model's response as a function of stimulus orientation and phase for $\lambda_1 = 10$. The input $x$ is filtered with a Gabor filter that has the same orientation preference as $w_0$. Note that this implementation differs from the hierarchical model of Hubel and Wiesel in that the input $x$ passes through a single linear filter, while in the standard hierarchical model $x$ passes in parallel to a number of phase shifted filters with the same orientation that in the second stage converge to a single filter. . . . 8

S5 **The model explains the relationship between stimulus contrast and phase sensitivity shown experimentally.** Modulation depth as a function of stimulus contrast, for different values of $\lambda_1$, and exponents $p$ and $q$, overlaid with experimental data taken from fig. 3c in (Meffin et al., 2015). . . . 9

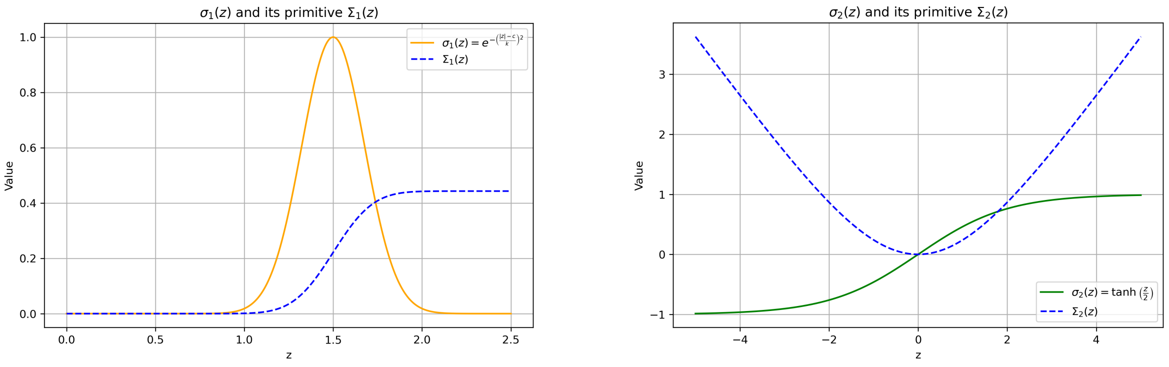


Figure S1: Graphs of the non-linearities $\sigma_1$ and $\sigma_2$ used in the experiments, together with the corresponding primitives $\Sigma_1$ and $\Sigma_2$

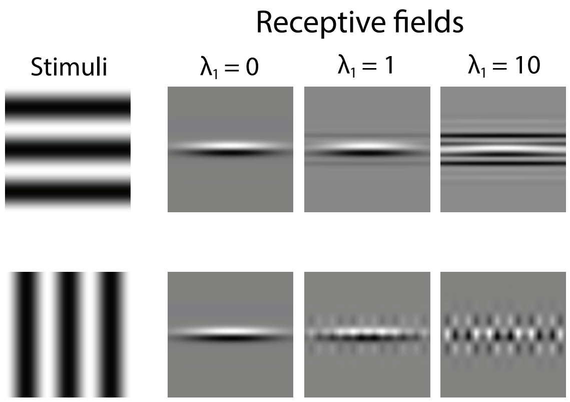


Figure S2: **The model has the same RF when its nonlinear terms are zero ($\lambda_1 = \lambda_2 = 0$) irrespective of the input stimulus, but for nonzero nonlinear contributions, its RF becomes stimulus dependent.** Results for a horizontal (top row) and a vertical (bottom row) sinusoidal grating stimulus (in all cases, $\lambda_2 = 0$). First column shows the stimuli, the other columns, the RFs for different $\lambda_1$ values. For $\lambda_1 = 0$, the RF does not depend on the stimulus and is the same in both cases, for $\lambda_1 \neq 0$ it is stimulus dependent.

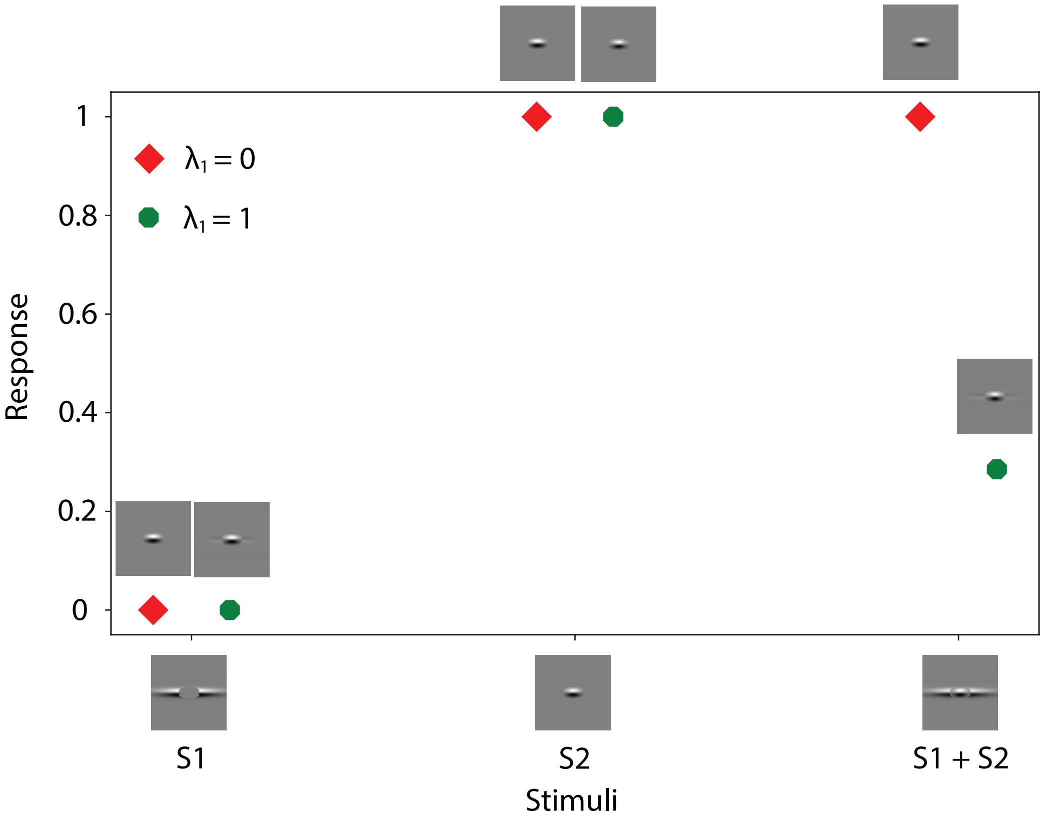


Figure S3: **For nonzero nonlinear contributions ($\lambda_1 = 1$), the model shows endstopping.** Normalized spiking response of the model (omitting the second nonlinear term, i.e. with $\lambda_2 = 0$) for three stimuli: $S1$ (part of a Gabor presented only on the sides, outside of the classical RF), $S2$ (a Gabor presented at the center, inside the classical RF), and their sum, $S1+S2$. The red diamond markers indicate the response of the unit when $\lambda_1 = 0$, and the green circle markers when $\lambda_1 = 1$. The images above the markers indicate the RF of the unit for the $\lambda_1$ value corresponding to the marker below and the stimulus corresponding to the abscissa position. For no nonlinear contribution ($\lambda_1 = 0$), the RF stays the same irrespective of the stimulus and the model does not show endstopping; the responses for $S2$ and $S1 + S2$ are the same. For a contribution from the nonlinear term ($\lambda_1 = 1$), the RF changes with the stimulus, and the response of the unit shows endstopping; it decreases for the elongated stimulus ($S1 + S2$) compared to the Gabor within the classical RF ($S2$), but does not respond to the side parts of the stimulus ($S1$) when they are presented in isolation.

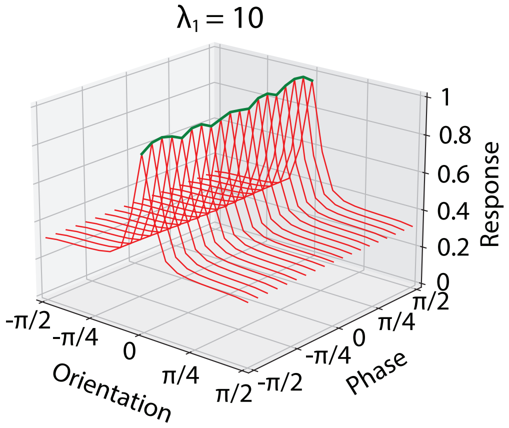


Figure S4: **The model produces a complex response with cortical input.** 3D plot of the model's response as a function of stimulus orientation and phase for $\lambda_1 = 10$. The input $x$ is filtered with a Gabor filter that has the same orientation preference as $w_0$. Note that this implementation differs from the hierarchical model of Hubel and Wiesel in that the input $x$ passes through a single linear filter, while in the standard hierarchical model $x$ passes in parallel to a number of phase shifted filters with the same orientation that in the second stage converge to a single filter.

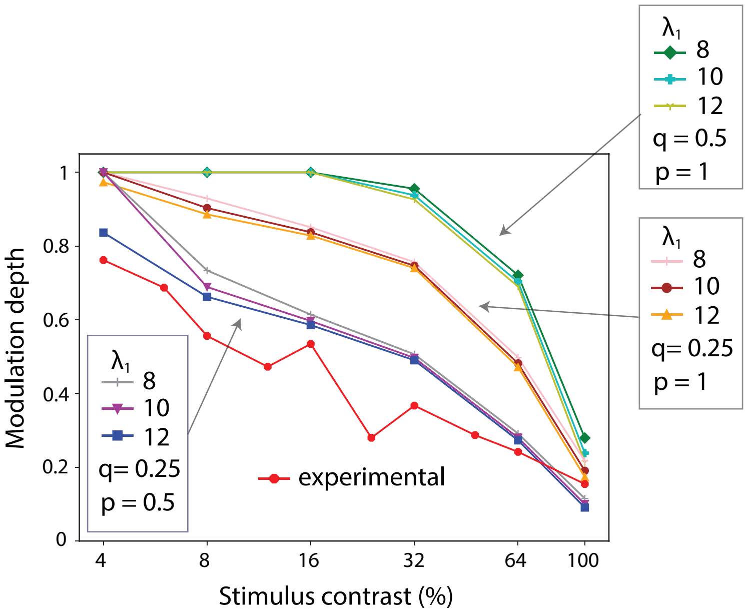


Figure S5: **The model explains the relationship between stimulus contrast and phase sensitivity shown experimentally.** Modulation depth as a function of stimulus contrast, for different values of $\lambda_1$, and exponents $p$ and $q$, overlaid with experimental data taken from fig. 3c in (Meffin et al., 2015).
